# Neuronal ensembles in visual cortex reactivate during hypnagogic sleep onset

**DOI:** 10.1101/2023.04.26.538480

**Authors:** Justin Lines, Rafael Yuste

**Author notes:** Correspondence should be addressed to J.L. new address: Dr. Justin Lines, PhD, Friedman Brain Institute, Dept. of Neuroscience, Icahn School of Medicine at Mount Sinai, S10 Hess Center for Science and Medicine, 1470 Madison Ave, New York, NY 10029.

## Abstract

During sleep onset (i.e. hypnagogia), visual hallucinations can occur in healthy human subjects following prolonged sensory experience. These percepts at sleep onset also occur in amnesiac patients and have signatures in human scalp electroencephalography recordings, suggesting a cortical origin. At the cellular level, neuronal ensembles, defined as groups of coactive neurons, dominate cortical activity and are causally related to perceptual states and behavior. To better understand the circuit mechanisms underlying hypnagogic visual hallucinations during sleep onset, we imaged neuronal activity in the rodent visual cortex during wake and sleep brain states following the repeated presentation of visual stimuli. We find that NREM1 sleep is the primary brain state at sleep onset, and neuronal ensemble properties during sleep are similar to wake. Interestingly, following prolonged visual stimulation, neuronal ensembles in visual cortex recruited by visual stimuli or locomotion had increased activity specifically during NREM1 sleep, consistent with the presence of hypnagogic hallucinations. Moreover, neuronal ensembles defined during visual stimulus periods were already present in visual cortex preceding visual stimulation. We propose that neuronal ensembles form an endogenous lexicon of cortical activity that are specifically recruited by sensory stimuli, and that specific neuronal ensembles become reactivated in visual cortex during hypnagogic sleep onset following prolonged sensory experience.

**One-Sentence Summary:** Prolonged sensory experience induces the activity of endogenous neuronal ensembles in rodent visual cortex during sleep onset, providing a circuit mechanism for the hypnagogic hallucinations reported in human subjects.

**Summary:** The encoding of perceptual memory is believed to rely on the coactivity of ensembles of neurons. Hypnagogic images during sleep onset have been reported in healthy human subjects following extended visual stimulus presentation. To investigate this phenomenon at the cellular scale, we monitored populations of neurons in mouse visual cortex with two-photon imaging across wake and sleep states following repeated visual stimulus presentation. We found that visually recruited neuronal ensembles had increased activity during brain states of sleep onset, matching human findings. Analyzing neuronal activity prior to visual stimulus presentation uncovered visually recruited ensembles were already present during wake and sleep states. We propose that hypnagogic images are encoded on a pre-configured lexicon of ensembles in the visual cortex.

## Introduction

Hypnagogia, the brain state occurring at sleep onset, where visual, auditory and somatosensory hallucinations can occur in healthy individuals ^1–4^, reported as images, sounds or tactile sensations. Following long sessions of playing Tetris, healthy subjects reported visual hallucinations of blocks during hypnagogia sleep onset brain states ^5^. In the same study, hypnagogic images were also reported by amnesiac patients ^5^, suggesting a non-hippocampal form of memory reactivation during sleep. More recently, functional magnetic imaging of human subjects has revealed neural forms of memory reactivation in visual cortex, and not hippocampus^6^, after repeated visual stimuli in offline rest periods ^6,7^. Here, we sought to determine the underlying neuronal code of hypnagogic images in the visual cortex during sleep onset.

At the cellular scale, perceptual states are believed to be encoded in organized ensembles of synchronous neuronal firing by small groups of neurons within the cortex ^8–13^. Interestingly, these neuronal ensembles can be activated by the exposure to a sensory stimulus but can also occur spontaneously, in the absence of any sensory or motor correlates ^14–16^. Previous studies have shown that spontaneous neuronal ensembles are statistically identical to visually recruited ensembles ^14–17^. This resemblance suggested the hypothesis that ensembles are the endogenous building blocks of cortical function that are recruited by sensory stimuli, from an internal configuration of patterns. Interestingly, rodents that experienced prolonged visual stimuli had increased cortical reactivation for tens of minutes afterwards ^18^. Therefore, another possibility is that sensory stimulus creates and “burns in” ensembles into cortical activity, and that these ensembles are later reactivated spontaneously as echoes of past sensory experience ^19^. To discern these possibilities, one potential clue comes from studies of sleep.

During sleep, the brain is placed in a quiescent state, disconnected from sensory input. Cortical activity during wake and rapid eye movement sleep (REM) displays high frequency brain waves ^20^. In contrast, cortical activity during non-REM (NREM) slow wave sleep is reduced to low frequency oscillations ^21^, where the newly formed patterns of neuronal activity are thought to spontaneously reactivate in a phenomenon known as reactivation ^22–25^. Indeed, neuronal activity has been shown to reactivate during wake, NREM and REM sleep ^23,25–28^. Memory reactivation is the spontaneous activation of newly formed neuronal ensembles, posited to improve the stability of ensemble activity and improve memory retention ^29–31^. The cortex is believed to act as long-term memory storage ^32^, where during sleep, communication between the hippocampus and cortex supports memory consolidation following learning ^33–35^. Previous reports have shown neuronal sequential reactivation during NREM sleep in the hippocampus and primary visual cortex during a spatial task ^23,25^. Molecular investigations of immediate early genes in visual cortex, and the modulation of their target neurons, uncovered sleep dependent mechanisms of visual specificity in contextual conditioning ^35^. Studies of memory reactivation have relied mainly on the learning of complex behaviors such as reward association or contextual encoding ^18,23,25,36^, with little work on the reactivation of neuronal ensembles encoding perceptual experience during sleep. The initiation of cortical memory reactivation events relating to specific brain states such as sleep onset (i.e. hypnagogia), is unknown.

Hypnagogic hallucinations found during sleep onset appear to represent a phenomenon of perceptual memory reactivation that is cortical in origin. The majority of cellular investigations on memory reactivation during sleep have been limited to the hippocampus following tasks of spatial learning and memory. Interestingly, hippocampal sequences of neuronal activity during a spatial task in animals naïve to the spatial context can be found during prior sleep states ^37,38^, suggesting sequences of memory traces ride atop a prebuilt organization in the hippocampus, and that this is likely due to an underlying organization of excitability ^37^. Cortex is a key component in theories on memory, and cellular studies of memory reactivation during sleep states, and the pre-allocation of neuronal ensembles in sensory cortices during sleep is needed to match the wealth of human studies ^7,32,39,40^.

Here we aimed to evaluate neuronal ensemble reactivation during hypnagogia in the visual cortex. We hypothesize that hypnagogic images will produce a signature at the cellular scale in the visual cortex during brain states of sleep onset following prolonged visual experience. Taking a conservative approach and presenting pseudorandom visual stimuli to avoid learning and the involvement of other brain regions, we test this hypothesis and further investigate the relation between visually recruited ensembles and ongoing cortical activity. Specifically, we imaged neuronal ensembles in the visual cortex during wake and sleep brain states before and after the presentation of a prolonged and pseudorandom visual stimuli in a head fixed mouse. First, we identify the brain state that wake generally enters sleep, then evaluate neuronal ensembles in the visual cortex across wake and sleep brain states. Next, we assess the spontaneous activity of visually recruited neuronal ensembles during the prior and subsequent wake and sleep stages to identify the brain state containing cellular signatures of memory reactivation. Our results are consistent with the hypothesis that sensory activity recruits pre-existing cortical neuronal ensembles that are then reactivated spontaneously, specifically during sleep onset (i.e. hypnagogic) brain states.

## Results

### NREM1 is the primary brain state at sleep onset

To record neuronal ensembles across wake and sleep states, we measured the activity of neuron populations using two-photon calcium imaging, in parallel with electrocorticogram (ECoG) and electromyography (EMG), in head-restrained mice (Fig. 1A-C). In addition to electrophysiology, an infrared camera was used to monitor the mouse to confirm sleep status and manually validate automatic sleep rating (Fig. 1B). We took an unbiased approach to rate sleep state, by implementing an automatic sleep rating algorithm to determine wake and sleep status of animals while recording neuronal calcium activity in layer 2/3 of the visual cortex (see Materials and Methods, Fig. 1C-D). Mice were handled for a week leading up to habituation on the head restraint system that took 2-5 days of consistent runs of 1-2 hours in length for mice to reliably sleep while being head restrained. Recordings were performed between Zeitgeber times (3-9), the room was temperature controlled to stay above 24° C, and a lavender scent was introduced. Following habituation to the experimental setup, calcium imaging recordings were classified and assigned to distinct behavioral states: locomotor, non-locomotor quiet wake, REM sleep and non-REM stages NREM1 and NREM2 sleep states (Fig. 1D-E). Collating the brain states found at sleep onset from wake throughout our recordings determined that NREM1 sleep was the primary sleep state during hypnagogia (Kruskal-Wallis: p < 0.001; quiet wake transition to: NREM1 sleep = 64.4 ± 11.0 % vs NREM2 sleep = 15.8 ± 7.5 % and REM sleep = 19.8 ± 9.2 %; p < 0.001; 47 experiments 12 animals: Fig. 1F), matching the sleep state found in humans during sleep onset ^41,42^. On average our recordings captured calcium activity from 202 ± 108 neurons in visual cortex per session (47 experiments, 12 animals). For each recording, calcium transients from individual neurons were analyzed to detect individual frames of spiking activity and create rasterized traces (Materials and Methods).

**Fig. 1.**
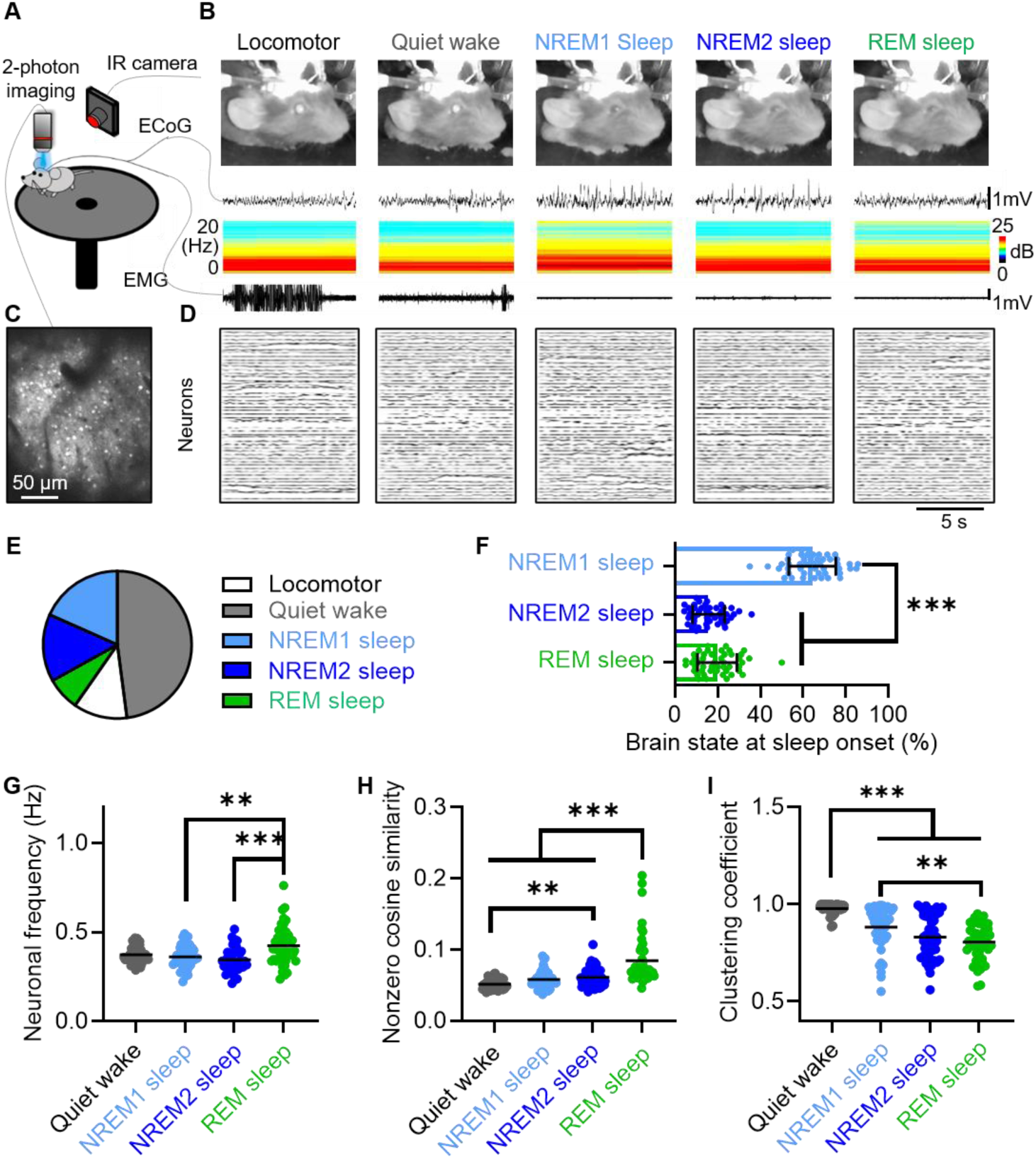
Imaging neuronal activity in visual cortex during head-restraint hypnagogia and sleep states. (**A**) Head-restrained imaging and electrophysiology. (**B**) Infrared imaging of mouse across wake and sleep states with ECoG traces, frequency spectrograms, EMG traces. (**C**) GCaMP6s expression in layer 2/3 of the primary visual cortex. (**D**) Neuronal Ca^2+^ traces from C across wake and sleep states. (**E**) Breakdown of brain states recorded during head restraint (Zeitgeber time 3-9). (**F**) Sleep state entered first from wake. (**G**) Frequency of neuronal calcium events. (**H**) Average nonzero cosine similarity between neurons across brain state. (**I**) Average clustering coefficient. Black lines denote mean. Dunn’s test after significant Kruskal-Wallis test. ** ≡ p<0.01, *** ≡ p<0.001

Since sleep has such a profound effect on population activity as seen in ECoG, we next asked what effect brain state had on cellular and network level neuronal activity in visual cortex. At the single neuron scale, we observed no change in neuronal activity between quiet wake and NREM sleep states, however we did observe a marked increase in neuronal frequency in REM sleep (Kruskal-Wallis: p < 0.001; 0.43 ± 0.10 Hz in REM sleep vs. 0.36 ± 0.06 Hz in NREM1 sleep, p < 0.01; and vs 0.34 ± 0.07 in NREM2 sleep; p < 0.001; 47 experiments; 12 mice; Fig. 1G), matching recent findings of increased firing of neurons in occipital regions during REM sleep ^43^. Next, we assessed properties of neuronal network activity across wake and sleep brain states. Cosine similarity represents moments of coactivity between neuron pairs, albeit not necessarily physically connected ^44^. The similarity between neuronal pairs is altered in models of neurocognitive disorders ^45^, and we asked whether this is also altered across brain state. We quantified the cosine similarity between two neurons as the overlap of active frames from rasterized traces normalized by each cell’s total firing (see Materials and Methods). When comparing neuron pairs with nonzero positive similarity, we observed network activity during REM sleep was increased compared to all other brain states (Kruskal-Wallis: p < 0.001; 8.4 ± 3.4 % in REM vs. 5.1 ± 0.7 % in wake, 5.8 ± 1.2 % in NREM1, 6.1 ± 1.3 % in NREM2: p < 0.001; 47 experiments, 12 animals: Fig. 1H). Similarity found during quiet wake was not significantly different than NREM1 sleep (p > 0.05) but was increased in NREM2 sleep (p < 0.01). To further investigate the functional network architecture, we quantified the average clustering coefficients of the network by creating adjacency matrices of connectivity, defined by positive values of cosine similarity. We found that the average clustering coefficient was reduced in all sleep states, when compared to quiet wake (Kruskal-Wallis: p < 0.001; 0.98 ± 0.03 in wake vs. 0.88 ± 0.11 in NREM1, 0.83 ± 0.11 in NREM2, 0.80 ± 0.09 in REM: p < 0.001; 47 experiments, 12 animals: Fig. 1I). In addition, clustering during REM sleep was significantly reduced compared to NREM1 sleep (p < 0.01). These results indicate that differences observed in the population level analysis of our calcium imaging aligns to the delineation of brain state we made from ECoG, and suggests processes during wake activate a denser cluster of neurons compared to sleep states void of sensory input. Finally, we show that NREM1 sleep, the primary state at sleep onset, shares some of the properties of neuronal activity found in the visual cortex during quiet wake.

### Neuronal ensemble properties during quiet wake are conserved in NREM sleep

Observing changes in the cellular and network activity suggested that encoding in the visual cortex may differ across brain state. One way that cortical circuits encode sensory information occurs via the synchronous firing of neurons forming neuronal ensembles ^12,16,46,47^. To explore this, we investigated the organization of neuronal ensembles found across wake and sleep states. We extracted neuronal ensembles from the rasters of neuronal activity across sessions where every brain state was observed using Non-Negative Matrix Factorization (NMF, see Materials and Methods) ^48,49^ (Fig. 2A-D). Briefly, we use NMF to reduce rasters of neuronal activity to clusters of synchronously active neuronal ensembles, with a minimum size of 4 neurons. Using this method, we detected neuronal ensembles occurring spontaneously across all wake and sleep brain states (Fig. 2E, Fig. S1). We detected on average 11.9 ± 7.6 ensembles per recording (Fig. S1C), and each ensemble detected consisted of 14.2 ± 4.6 neurons (Fig. S1D). Almost all detected ensembles were present in quiet wake (99.1 ± 2.6 %), and significantly fewer in sleep states (Kruskal-Wallis: p < 0.001; vs. 91.7 ± 12.1 % in NREM1: p < 0.01, vs. 88.8 ± 14.2 % in NREM2: p < 0.001, vs. 74.9 ± 21.7 % in REM sleep: p < 0.001; 47 experiments, 12 animals: Fig. S1E). Together, this result suggests that most neuronal ensembles become active spontaneously across all brain states, with a majority present during wake.

**Fig. 2.**
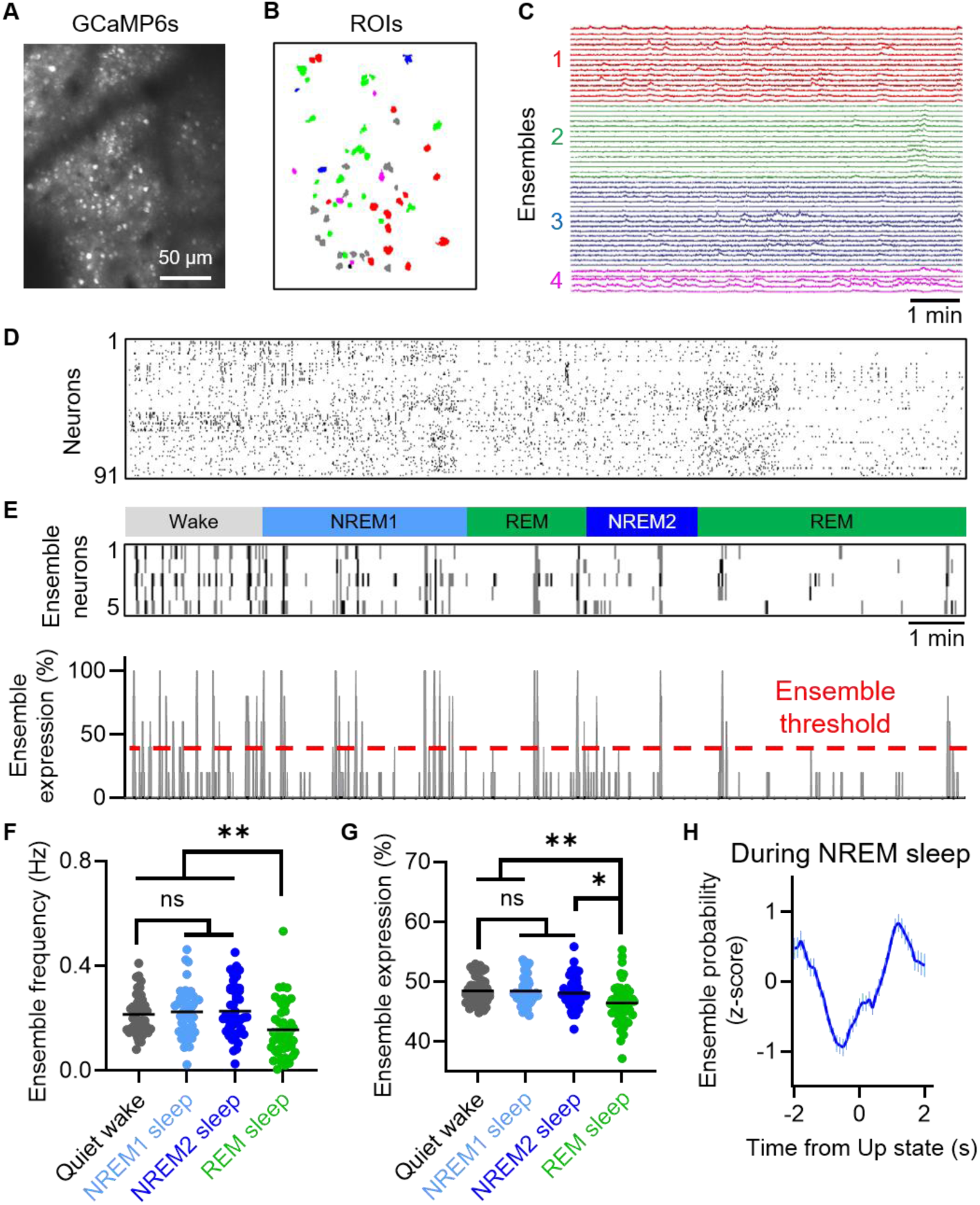
Neuronal ensembles present during NREM1 sleep have similar properties as in wake. (**A**) GCaMP6s positive neurons. (**B**) NMF extracted ensembles overlayed on ROIs. (**C**) Traces from the first four ensembles from like colored cells in B. (**D**) Sorted raster of neuronal activity based on NMF labeling. (**E**) Raster of cells within ensemble 4 in C across wake and sleep (top), and its expression of proportion of cells active (bottom) with 35% ensemble threshold. (**F**) Ensemble frequency. (**G**) Ensemble expression. (**H**) The probability of ensemble activity related to UP state activity during NREM sleep. Dunn’s test after significant Kruskal-Wallis test. * ≡ p<0.05, ** ≡ p<0.01

To characterize the properties of neuronal ensembles across brain states, we introduced the metric of ensemble expression, to capture the extent of activation for an individual ensemble over time. Ensemble expression is defined as the percentage of neurons synchronously active within an individual ensemble at any given time (Fig. 2E), e.g. if 4 neurons are synchronously active from an ensemble comprised of 10 neurons, then the ensemble expression at that frame would be 40%. Next, we defined that an ensemble was active if it had an expression of over three standard deviations from the mean expression from all neuronal ensembles identified (5.9 ± 9.4% ensemble expression; > 35%; Fig. S1A-B), encompassing values from the long tail of the distribution of ensemble expression. Using this definition, we quantified the number of times an ensemble is active per second, termed the ensemble frequency. In contrast to our previous result of increased single neuron firing frequency during REM sleep (Fig. 1G), we found that REM sleep had the lowest neuronal ensemble frequency compared to quiet wake and NREM sleep (Kruskal-Wallis: p < 0.001; 0.16 ± 0.11 Hz in REM sleep vs. 0.21 ± 0.07 Hz in quiet wake, 0.22 ± 0.09 Hz in NREM1 sleep, 0.23 ± 0.10 HZ in NREM2 sleep: p < 0.001; 47 experiments, 12 animals: Fig. 2F). At the same time, the neuronal ensemble expression, quantified as the percentage of neurons comprising an ensemble activation, was significantly decreased during REM sleep compared to wake and NREM sleep (Kruskal-Wallis: p < 0.01; 46.5 ± 3.5 % in REM sleep vs. 48.5 ± 2.2 % in wake: p < 0.01; vs. 48.5 ± 2.4 % in NREM1 sleep: p < 0.01; vs. 48.1 ± 2.6 % in NREM2 sleep: p < 0.05; 47 experiments, 12 animals: Fig. 2G). Similarly to single cell analysis, there was no significant difference between quiet wake and NREM sleep for both ensemble frequency (p > 0.05) and ensemble expression (p > 0.05). Together, these results demonstrate that spontaneous neuronal ensembles are present in all wake and sleep states. Interestingly, results showing heightened single cell neuronal activity during REM sleep did not translate to ensemble activity, being rarer and less pronounced specifically in REM sleep, demonstrating the importance and novelty in using these analyses. Interestingly, the properties of spontaneous neuronal ensembles during wake are also not significantly different to NREM sleep, suggesting neuronal ensembles act similarly across these brain states.

One prominent electrical feature in the ECoG during slow-wave NREM sleep is a cortical UP state, where the cortex switches from low to high levels of network activity, which are believed to organize spontaneous firing of cortical activity ^20,23,33,50^. We next asked whether the neuronal ensembles we identified from calcium imaging were modulated by UP states occurring within the simultaneously recorded ECoG signal. UP states were detected during NREM sleep brain states from a filtered ECoG trace (Fig. S2, see Materials and Methods). UP states within NREM sleep occurred at a frequency of 0.68 ± 0.29 Hz, in agreement with previous reports using similar methods ^50^. When we measured nearby detected UP states, we observed that neuronal ensemble probability was reduced preceding UP states and increased through the passage of the UP state in the cortical circuit (Fig. 2H). These findings match recent work analyzing single cell neuronal activity during UP states using a similar approach ^50^, illustrating UP states as transitions of inhibition leading into heightened neuronal activity, suggesting neuronal ensembles are actively being processed within NREM sleep brain states.

### Repeated visual stimulation increases the reactivation of task specific ensembles during subsequent NREM1 sleep

Hypnagogic images are believed to occur at sleep onset following repeated visual stimulus viewing ^5,6,42^. To explore this in cellular detail, we evaluated neuronal activity in visual cortex across wake and sleep states, before and after the presentation of repeated drifting visual gratings (Fig. 3A-D). Gratings were presented for 2 s with 3 s interstimulus interval and were repeated in a pseudorandom order in four directions (up, down, left, right) over a ten-minute-long stimulation block. Visual stimuli were presented in pseudorandom order to avoid habituation and novelty, which affect spontaneous ensemble properties ^51,52^. Rest periods were recorded during the 30 minutes directly before and after visual stimulus where animals could sleep. We did not find any significant changes in the similarity or clustering coefficients after visual stimulus presentation, suggesting that this type of visual stimulation does not alter the functional connectivity of wake and sleep states in the visual cortex. We also did not observe a change in UP state frequency during NREM sleep states after visual stimulus.

**Fig. 3.**
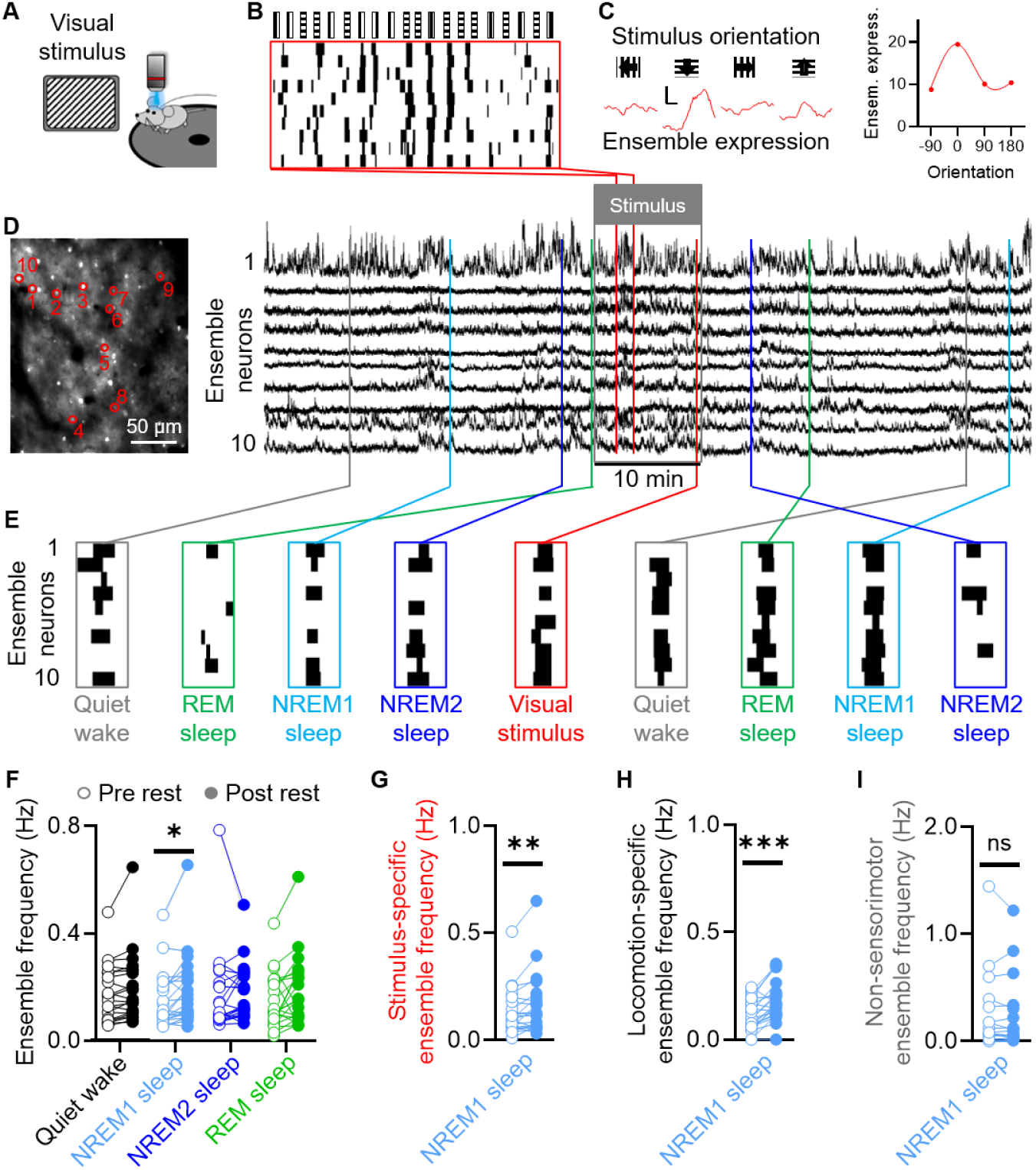
Prolonged visual stimulation increases ensemble activity in V1 during NREM1 sleep. (**A**) Scheme to present repetitive visual stimulus. (**B**) Raster of an ensemble with visual presentation. (**C**) Average visually evoked expression (left) and tuning curve (right) of ensemble in B. Scale is 5% and 1 s. (**D**) GCaMP6s positive neurons with red outlines over ensemble neurons (left) and traces (right) from B. (**E**) Frames of high neuronal ensemble expression in each brain state. (**F**) Change in ensemble frequency after stimulus based on brain state. Change in ensemble frequency during NREM1 sleep in stimulus-specific (**G**), locomotion-specific (**H**), and non-sensorimotor ensembles (**I**). Bonferroni corrected Wilcoxon test after significant Friedman test. * ≡ p<0.05, Wilcoxon test. ** ≡ p<0.01, *** ≡ p<0.001

We then turned our attention to ensembles and their presence during sleep states before and after visual stimulus presentation. We labeled ensembles using NMF constrained to frames during the stimulus block and then monitored the presence of task specific ensembles during the rest periods before and after visual stimulus. We detected 13.0 ± 9.1 ensembles per recording in this manner, which was similar to the number of detected ensembles found in spontaneous recordings (Fig. S1C). Surprisingly, we found neuronal ensembles defined during the stimulus period were also present across brain states after, as well as before, visual stimulus (Fig. 3D-E), suggesting the presence of ensemble reactivation as well as pre-existing cortical activity motifs respectively. Specifically, of ensembles defined in the stimulus period, 98.7 ± 2.8 % were present in the pre rest period, and 98.8 ± 3.3 % of these ensembles were also found in the post rest period. These results suggest that neuronal ensembles defined during the stimulus period in the visual cortex are present before, during and after visual stimulus presentation.

We then investigated if the visual stimulus presentation could alter the amount of ensembles present spontaneously in different wake and sleep states. Again, to quantify neuronal ensemble properties, we defined ensembles using NMF only on frames during the stimulus period, to determine the frequency of these ensembles in individual brain states, during resting periods before and after visual stimulus presentation. Comparing across brain state, we observed no significant difference in ensemble expression before and after visual stimulus (p > 0.05). Interestingly, when testing for the frequency of spontaneous neuronal ensembles, we found an increase following visual stimulation, specifically during NREM1 sleep (Friedman: p < 0.05; ensemble frequency during NREM1 sleep at 0.15 ± 0.10 Hz before vs. 0.19 ± 0.13 Hz after; p < 0.05 Wilcoxon test with Bonferroni correction; 22 experiments, 12 animals: Fig. 3F). At the same time, we did not observe changes in neuronal ensemble frequency in quiet wake, REM sleep or NREM2 sleep following visual stimulation (p > 0.05). Taken together, repeated visual stimulus presentation increases the reactivation of spontaneous neuronal ensemble activity specifically during NREM1 sleep, the sleep stage we found as the primary brain state of sleep onset matching human findings ^5,42^, possibly uncovering the cellular underpinnings of hypnagogic images during sleep onset following prolonged visual stimulus.

### Specific visual or locomotor neuronal ensemble have increased reactivation during NREM1 sleep

Identifying neuronal ensembles only during the stimulus period biased the detection of ensembles encoding visual stimulus, however some ensembles appeared to be unrelated to visual stimuli, e.g. locomotor. To address this, we categorized our collection of detected ensembles to evaluate the reactivation of ensembles in an encoding-specific manner. Ensembles were classified into three main groups: stimulus-specific, locomotor-specific, and non-sensorimotor, based on their significant positive Pearson’s correlation of ensemble activation to the stimulus presentation, locomotor or neither respectively (see Materials and Methods). Throughout our recordings, we detected 44.7% as stimulus-specific ensembles, 37.2% as locomotor-specific, and 18.1% as non-sensorimotor ensembles (Fig. S3). Closer examination into the content of non-sensorimotor ensembles showed them to be comprised of 34.6% as anti-correlated to sensory stimulus, 15.4% trending toward stimulus correlation, and 50.0% correlated to the delta brain rhythms, most likely related to periods where the mouse was not attending (Fig. S3).

Based on our finding that ensembles defined during periods of visual stimulus are increased during NREM1 sleep states, we next asked if this increase in ensemble activity following visual stimulus was constrained to ensembles encoding for specific elements of the mouse experience. First, we found that the frequency of visual stimulus-specific ensemble activity was significantly increased during NREM1 sleep following visual stimulus presentation (0.13 ± 0.11 Hz before vs. 0.16 ± 0.14 Hz after: p < 0.01; 22 experiments, 12 animals: Fig. 3G). Next, we assessed ensembles defined during the visual stimulus period that were significantly correlated to locomotor behavior and observed that locomotor-specific ensembles were also significantly increased during NREM1 sleep following visual stimulus presentation compared to before (0.11 ± 0.07 Hz before vs. 0.18 ± 0.09 HZ after: p < 0.001; 22 experiments, 12 animals: Fig. 3H). In contrast, ensembles that were neither correlated to visual stimulus presentation nor locomotor, which we call non-sensorimotor ensembles, had no significant change in ensemble frequency during NREM1 sleep before compared to after visual stimulus (0.20 ± 0.34 Hz before vs. 0.19 ± 0.32 Hz after: p > 0.05; 22 experiments, 12 animals: Fig. 3I). Taken together, these results demonstrate that the increases we observed in ensemble frequency within NREM1 sleep following a visual stimulus presentation in a head-restrained system was specific to ensembles that were recruited by either the visual stimulus or locomotor activity. On the other hand, ensembles that were not related to either sensorimotor dimension showed no frequency changes within NREM1 sleep, suggesting that ensembles encoding for aspects outside the behavior, while still part of the underlying architecture and spontaneous activity in offline brain states, do not experience a boost in processing during subsequent NREM1 sleep as behavior-specific neuronal ensembles do. This matches findings in human subjects that hypnagogic hallucinations can include visual as well as somatosensory perceptions during sleep onset ^42^.

### Visually recruited neuronal ensembles are present in visual cortex before visual stimulation

Next, we wanted to follow up on the neuronal ensemble activity we observed during rest periods prior to visual stimulation. Importantly, neuronal ensembles were present before stimuli presentation in experiments even where mice were naïve to visual stimulus (Fig. 4A), that would later go on to form the neuronal activity underlying responses in the visual cortex to visual stimuli (Fig. 4B). Overall, neuronal expression in the rest period prior to stimulus presentation was not different between experiments where mice were naïve to the stimulus or not (p < 0.05), and this was also true for ensemble frequency (p < 0.05). We compared the stimulus defined ensemble frequency during the stimulus and non-locomotor rest periods and found a significant change across these time periods (0.17 ± 0.11 Hz pre-stimulus, 0.52 ± 0.21 Hz during stimulus, 0.19 ± 0.12 Hz post-stimulus: Kruskal-Wallis: p < 0.001; 22 experiments, 12 animals: Fig. 4C). Comparing individual periods, we found that ensemble frequency during stimulus presentation was significantly higher than during rest periods (p < 0.001), as to be expected in the visual cortex when providing visual stimuli.

**Fig. 4.**
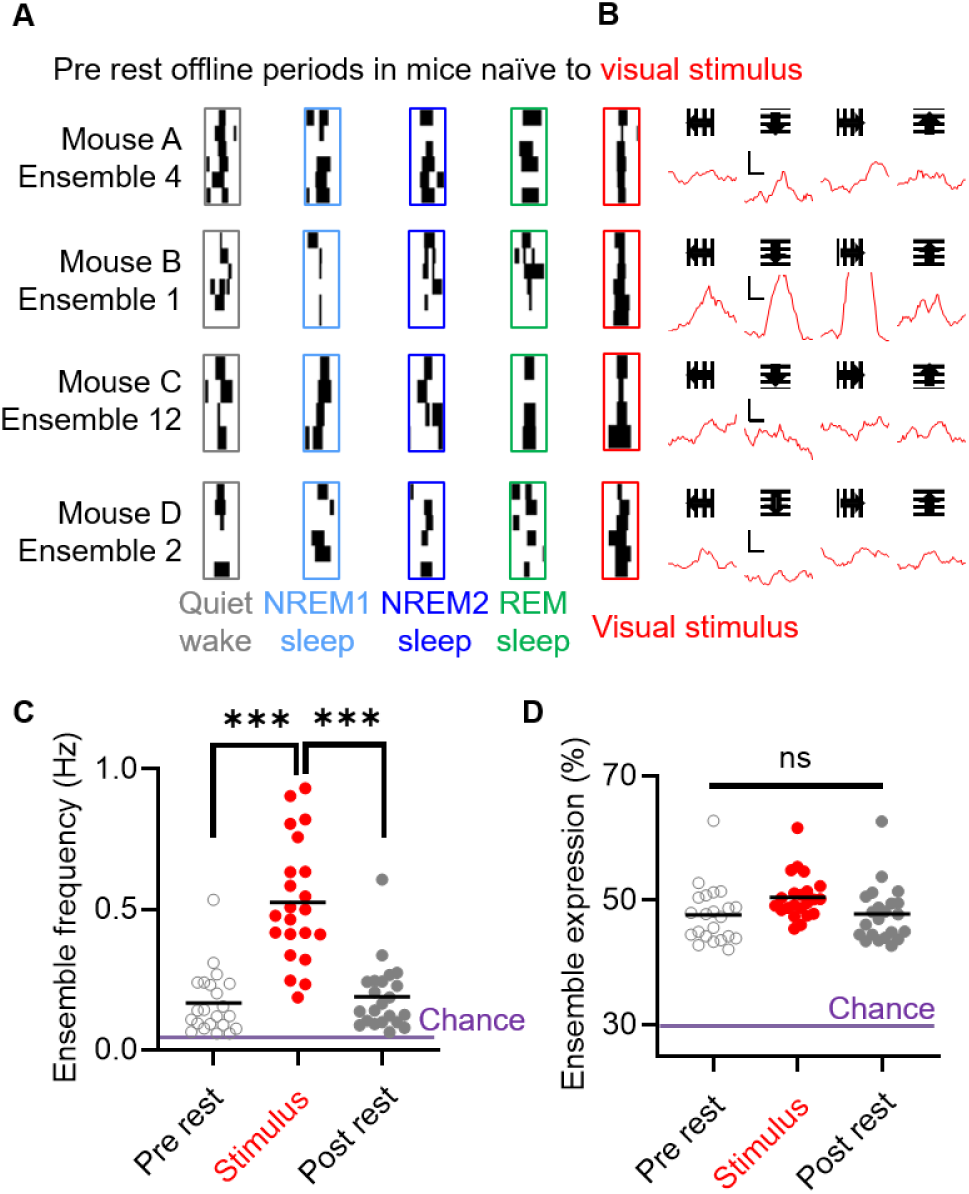
Visual stimuli recruit pre allocated neuronal ensembles in visual cortex already present in pre-stimuli offline brain states. (**A**) Visual recruited neuronal ensembles in naïve mice that were present during the rest period preceding initial visual stimulus in B. (**B**) Average neuronal expression from neuronal ensembles during the initial visual presentation. (**C**) Frequency of stimulus (red) defined ensembles during pre (gray) and post (filled) non locomotor rest periods compared to shuffled defined chance (purple). (**D**) Ensemble expression of stimulus (red) defined ensembles across pre rest (gray) and post rest (filled) periods compared to shuffle (purple). Dunn’s test after significant Kruskal-Wallis test. *** ≡ p<0.001

To confirm that neuronal ensemble activity during rest periods is not due to chance, we compared our experimentally defined neuronal ensemble frequency during non-locomotor rest periods with the frequency found in a distribution of these rasters shuffled. We created a shuffled distribution of chance for neuronal ensemble frequency by identifying ensembles present in a circular shuffled raster, 10,000 times per raster. Indeed, the frequency of neuronal ensembles present during offline states prior to visual stimulus was significantly higher than found in a shuffled distribution (0.17 ± 0.11 Hz experimental vs. 0.05 ± 0.06 Hz shuffled; p < 0.001: Fig. 4C), and this was similarly true for post stimulus rest periods (0.19 ± 0.12 Hz experimental vs. 0.05 ± 0.04 Hz shuffled; p < 0.001: Fig. 4C). These results suggest that neuronal ensembles in the visual cortex are present before, during and after visual stimulus presentation yet they are more active during the presentation of a visual stimulus. Across all recordings, average ensemble expression was not significantly different between the stimulus period and rest periods (Kruskal-Wallis: p > 0.05; 47.8 ± 4.5 % pre-stimulus vs. 50.5 ± 3.6 % during stimulus, and 48.3 ± 4.0 % post-stimulus; 22 experiments, 12 animals: Fig. 4D), suggesting that visually recruited ensembles are similarly expressed whether active during the stimulus period or spontaneously active during rest. As above for ensemble frequency, we compared average ensemble expression to a shuffled distribution and found that ensemble expression during pre rest periods was significantly higher than shuffled (47.8 ± 4.5 % experimental vs. 28.7 ± 11 % shuffled; p < 0.001; 22 experiments, 12 animals: Fig. 4D), and this was true for post rest periods as well (48.3 ± 4.0 % experimental vs. 30.0 ± 12 % shuffled; p < 0.001; 22 experiments; 12 animals: Fig. 4D). Note that shuffled data had an ensemble expression below our threshold for ensemble detection, due to the occurrence of many ensembles not being present in shuffled datasets. Further, compared to shuffled distributions, our data supports the hypothesis that visual stimuli recruit neuronal ensembles that are already present in the visual cortex.

## Discussion

Hypnagogic images have been reported in healthy human subjects after prolonged visual experience ^5,42^, and we show this effect at the level of neuronal ensembles in the visual cortex. Unlike human subjects that can report the perceptual experience of visual hallucinations at sleep onset ^42^, we must rely on physiological measures of cellular activity. We observe groups of synchronously firing neurons that are recruited by repeated visual stimuli and undergo a significant increase in activity during subsequent states of sleep onset (i.e. NREM1). Note that repetitive visual stimuli increased neuronal ensemble reactivation only during NREM1 sleep, yet these neuronal ensembles were already present across all brain states preceding visual stimuli, supporting the hypothesis that hypnagogic images, and stimulus recruited neuronal ensembles, come from a pre-allocated internal lexicon of network encodings in the visual cortex.

We found that NREM1 was the primary sleep state at sleep onset, and this matches the brain state where human subjects report visuomotor hypnagogic hallucinations ^42^. This specific stage of sleep may place the cortex in a unique environment. NREM1 sleep is a quiescent state void of sensory experience and without major neuromodulatory control, such as wake, REM sleep, and NREM2 sleep driven largely by acetylcholine, serotonin and norepinephrine ^53^. Additionally, during NREM1 sleep, the cortex may receive less input from thalamic and limbic brain areas heavily involved during wake and deeper sleep states resulting in more self-generated cortical activity. NREM1 sleep may be when the cortex is in a unique neuromodulatory environment with reduced inputs from other brain regions to produce local self-generated neuronal activity, and represent a key initializing step in offline processing during sleep ^41^.

The effect of sleep could be observed in cortical networks in the parameters of cosine similarity and clustering. During sleep states, compared to wake, the increase in cosine similarity of nonzero pairs came with a decrease in average clustering, suggesting network states of heightened similarity in firing patterns found in REM and NREM sleep. This matches NREM and REM sleep states being widely understood to be synchronous states compared to desynchronized waking states ^21,53^, dominated by oscillations in the local field potential ^20^. In addition, reports have found differential neuronal activity between wake, REM and NREM sleep ^54–57^, and changes in functional network architecture at the mesoscopic scale of motor cortices across wake and sleep ^58,59^.

The encoding of percepts on cortical neuronal networks is believed to occur by neuronal ensembles during the repeatable cofiring of neurons ^11,12,16,46,47^. Indeed, using NMF we discovered that neuronal ensembles were active across all vigilance states and that ensembles present during REM sleep had reduced expression and frequency of occurrence comparatively. This reduction in neuronal ensemble quality during REM sleep was in contrast to our findings that single neurons had increased activity specifically during REM and demonstrates the importance of the analyses used here. Further, this was in line with previous accounts of REM sleep being associated with distinct cortical dynamics, where activity is higher and less organized ^43^. Interestingly, properties of neuronal ensembles were no different between wake and NREM sleep phases, suggesting a conserved encoding between these two brain states.

Prolonged pseudorandom visual stimulation increased reactivation of neuronal ensembles specifically in NREM1 sleep. Human subjects describe hypnagogic hallucinations during sleep onset that are specific to the sensory experience before sleep ^5^, characterized as NREM1 sleep stages ^42^, and functional magnetic imaging confirms these activities in visual cortex ^6,7^. Indeed, we show that neuronal ensembles in visual cortex that were recruited by visual stimuli have increased activity during NREM1 sleep stages following prolonged visual stimulation. When accounting for the specific content of individual neuronal ensembles, we observed that the spontaneous reactivation during NREM1 were content-specific. Interestingly, the main factors that the mouse is exposed to in the paradigm were visual stimulus and locomotion on the disk, and the ensembles encoding this behavior in visual cortex are preferentially reactivated over ensembles that are less related to the task. This adds to animal work outlining increases in spontaneous neural activity following stimulation ^18,19^, and the role of ensemble reactivation in visual cortex during sleep following contextual encoding ^23,35^. Previous studies have focused on the presence of memory reactivation following learning and the role of cortex along with limbic brain regions to improve memory. It is unknown the effect of hypnagogic hallucinations on cortical circuits; whether no effect, positive or negative. It may be that cortical reactivations during this brain state are a mechanism of forgetting ^60^, to enhance these signals in order to eventually normalize them to the rest of the network ^18^.

Neuronal ensembles detected during the visual stimulus presentation were also found in spontaneous activity preceding the stimulus period. These results suggest that the neuronal ensemble activity recruited by visual stimuli are already present in the visual cortex. Sensory stimulus can recruit previously encoded neuronal ensembles in primary visual cortex ^16^, and here we find this translates across sleep states as well. Studies have demonstrated that neuronal sequences are pre-allocated in the hippocampus ^37,38^, and our findings suggest neuronal ensembles are also pre-allocated in cortical networks, providing the cellular level assessment adding to human studies ^39,40^. Altogether, this suggests that network responses to visual stimulus are not newly formed ensemble activity, but that, instead, the cortex is recalling an ensemble, as a circuit module, that is already present in the network.

Taken together, our results provide a bridge between human and rodent studies of memory reactivation following visual stimulation that explains the similarity between neuronal ensembles in spontaneous and visually recruited activity by monitoring brain state along with cellular activity. Rather than the original hypothesis that cortical activity during sensory stimulation directly generates neuronal ensembles, our data suggests that visual stimulation recruits endogenous neuronal ensembles in the visual cortex. By repeated applications, these neuronal ensembles then have increased reactivation at sleep onset. Overall, our data indicate that visually recruited neuronal ensembles in cortical circuits relate to rigid pre-existing neuronal ensembles, and yet react to sensory stimuli with enhancement of reactivation during NREM1 sleep onset.

## Materials and Methods

### Experimental Design

The main objective of the study was to interrogate the properties of neuronal ensembles in the visual cortex during sleep onset, wake and sleep brain states, specifically before and after the presentation of a visual stimulus. In order to achieve this overarching objective, mice were head-restrained to control their field of view as well as allow the continual imaging of visual cortex across wake and sleep states. Mice were transgenic that expressed a calcium indicator in excitatory neuronal populations, and were implanted with tungsten wire to record electrocorticogram and label brain states of wake and sleep.

### Proper animal use and care

All the procedures for handling and sacrificing animals were approved by the Columbia University Institutional Animal Care and Use Committee (IACUC) in compliance with the National Institutes of Health guidelines for the care and use of laboratory animals. We used 12 mice, both female and male transgenic animals (*VGluT1*-GCaMP6s) that were 2-6 months of age, kept on a continuous 12h light/dark cycle and freely available to food and water. Mice were regularly monitored for health and welfare by trained animal technicians and veterinarian staff.

### Stereotaxic surgery for cranial window, ECoG and EMG

Mice were anesthetized with 1.5-2% isoflurane and placed in a stereotaxic atop a heating pad with a probe to maintain body temperature at 37° C, and faux tears were applied to prevent corneal dehydration. Carprofen (5 mg/kg), Dexamethasone (0.6 mg/kg), and Enrofloxacin (5 mg/kg) were administered subcutaneously. Hair was removed from the scalp using Nair Hair Removal before ethanol and chlorhexidine was applied. Lidocaine was administered locally to the scalp followed by an incision made down the midline of the scalp to expose the skull. A hole was made over the primary visual cortex ^61^ (V1; in mm from bregma: -3.4_a-p_, 2.1_m-l_) and a tungsten electrode was placed atop the exposed cortex to act as the lead for recording electrocorticogram (ECoG). Next, an anchor screw soldered to a wire used as the ECoG reference was placed over the right frontal plate. Two wires were placed in the nuchal muscle to record electromyography (EMG). Next a titanium headplate was centered over the contralateral visual cortex and cemented onto the exposed skull using dental cement. After the cement dried, a 3 mm craniotomy was made over the contralateral primary visual cortex (centered in mm from bregma: -3.4_a-p_, -2.1_m-l_), and the dura was removed. Finally, a 3 mm glass coverslip was placed on the exposed cortex and fixed using super glue and dental cement. Mice recovered in their homecage and were provided with postoperative care that included regularly monitoring for signs of pain and distress as well as Carprofen (5 mg/kg), Dexamethasone (0.6 mg/kg), and Enrofloxacin (5 mg/kg) administered subcutaneously over three days. Animals were left to recover for 1-2 weeks before experiments were performed.

### Animal habituation to head restraint and experimental design

During the first week, mice were handled by the experimenter during the day and in the room where imaging would be performed. Following the week of handling, mice began habituation to the head restraint system. The head restraint system was comprised of construction rods connected to the imaging breadboard and bolted onto the titanium headplate fixed to the animal’s head, where the mouse was able to freely run on a disk attached below (Fig. 1A). The initial habituation to the head restraint system lasted 5 minutes, with subsequent days of head restraint time doubling until session length was at a maximum of 2 hours. Mice were habituated to the head restraint system until they were reliably sleeping. To aid sleep, the imaging room was kept above 24° C and a lavender scent was added. Following habituation, after mice were consistently sleeping across sessions, we performed simultaneous imaging and electrophysiology in spontaneous recordings for 1-2 hours in length, where mice were free to run or sleep. After spontaneous recordings were completed, mice were habituated to the monitor that would be used to show visual gratings for 1-2 sessions before experimental recordings. Imaging sessions were discarded if the animal did not sleep and/or under distress.

### In vivo two-photon calcium fluorescence imaging

Above the head mounted animal was an objective for in vivo imaging of the exposed mouse cortex, performed using a custom two-photon system comprised of Ti:sapphire lasers (MaiTai DeepSee at 920 nm). Imaging was done through optical gel under a 25x water immersion objective lowered over the center of the glass coverslip to image visual cortex with the focal plane at 100 – 300 µm below the cortical surface to record from layers 2/3. The microscope captured 512×512 digital images (500 µm^3^) at 31 Hz via a resonance galvanometer. Data included in this study were from videos obtained for 1-2 hours.

### Electrophysiological recordings

Electrocorticogram (ECoG) was recorded using an A-M Systems Model 3000 at a bandpass filter of 1 Hz – 1 kHz. Electromyography (EMG) was recorded using a Warner Instruments DP-301 with a bandpass filter of 1-100 Hz. Locomotion signal was recorded using a light detector through spokes on the disk edge the mouse ran on, that translated light passage from disk spokes to a voltage. ECoG, EMG, locomotion and stimulus presentation were sampled at 5 kHz and fed into the imaging PC via a National Instruments BNC_2110 digital acquisition system. Imaging and electrophysiology data types were read and synchronized in time through a PC running the program MScan, capturing both mediums simultaneously.

### Sleep automatic rating

Sleep rating was done semi-automatically using ECoG, EMG and locomotion signal as in ^62–64^. The ECoG frequency content and EMG activity was monitored to determine brain state based on number of standard deviations (SD) over the mean. The EMG signal was high-pass filtered at 25 Hz. The ECoG signal was low-pass filtered below 250 Hz and individual frequency bands were quantified from a short-time Fourier transform into delta (1-4 Hz) and theta (6-10 Hz) frequency bands. Sleep states were constrained to periods where locomotion was below 0.3 SD over the mean. All times above 0.3 SD from the locomotion signal were denoted as locomotion state. From the remaining periods of non-locomotive time, individual sleep brain states were defined based on the following levels of the EMG signal and ECoG spectra: below 1 SD EMG and over 0.5 or 1 SD delta activity (1-4 Hz) denote NREM1 or NREM2 sleep respectively, below 0.5 SD EMG with over 1.5 SD theta (6-10 Hz) divided by delta activity denotes REM sleep. Only epochs that lasted at least 15 seconds were accepted. All other time periods were counted as quiet wake. All automatic rating of wake and sleep states were verified with visual assessment of the ECoG and EMG activity along with visual confirmation of animal behavior from an infrared camera recording of mouse behavior.

### Up state detection

Within NREM slow-wave sleep, cortical activity switches between low and high network activity with the slow passing of cortical Down and Up states respectively ^21,65,66^. Up states, defined as periods of heightened network activity were detected in NREM sleep brain states of band pass filtered ECoG traces between 5 and 250 Hz. Up states were labeled if the filtered ECoG trace reached 3 standard deviations from the mean. Up state detection was manually checked to observe the characteristic deflection and high frequency activity in the raw ECoG trace ^50,62^.

### Visual stimulation

Drifting Gabor patches were presented to the visual field of the mouse on a monitor connected to a PC running Psychtoolbox ^67^ (psychtoolbox.org). Custom MATLAB code using Psychtoolbox was written to present Gabor patches along four directions (up, down, left, right) in pseudorandom order for 2 s in duration with 3 s interstimulus times of black screen and repeated 120 times over a 10-minute span. Rest periods of 30 minutes were also recorded before and after the stimulus presentation.

### Calcium image processing

To reduce computational time, images were frame-averaged by 3 to create videos at 10.3 Hz. All images of a recording were corrected for translational motion by registering to an average image created from the middle 5 minutes of the recording using TurboReg, a plugin for ImageJ. Next, regions of interest (ROIs) were created automatically from thresholding correlation images created from 10-minute imaging blocks and later combined to create ROI maps that were then updated manually using custom MATLAB scripts. Fluorescent traces were created by averaging over pixel values within ROIs as in ^16^. Fluorescent calcium traces from individual neurons were then binarized with an event detection as in ^45^ and compiled into rasters for each recording.

### Population and network measures of activity

To analyze population activity across brain state, cosine similarity between a neuron pair was analyzed by taking the cosine similarity of vectors of rasterized activity as in ^45^. Cosine similarity defined in this manner is constrained to values between 0 and 1 for no overlap and fully overlapped respectively. For individual brain states, only frames that included that specific state were used to quantify cosine similarity. Clustering coefficients were quantified by taking all nonzero cosine similarity matrices as adjacency matrices as input into the MATLAB function clustCoeff() ^68^.

### Neuronal ensemble detection using Non-negative Matrix Factorization

Neuronal ensembles were detected from rasterized traces using Non-negative Matrix Factorization (NMF) ^48,49^. NMF is a numerical clustering algorithm that was used to decompose an individual raster into two positive valued arrays: one composed of values representing the neuronal inclusion within each ensemble, and an array representing the ensemble weight at every frame. NMF requires an input of the number of components (i.e. the number of ensembles), and we determined the number of components in each raster by performing a principal component analysis on the raster, and finding the number of principal components with variances that exceeded 95% of the variance explained from a PCA performed on a shuffled raster as in ^69^. Shuffling was performed on individual rows of the raster by block shuffling the time series 10,000 times with blocks of random length, in order to preserve neuron firing rate. We then input the number of components found into the NMF function along with the original raster. NMF was run using the Alternating Least Squares algorithm ^48^. To ensure the stability of the solution, the NMF function was run with a tolerance of 10^-6^ at a max of 100 iterations, and we performed 100 replicates with random initial conditions. From the output array of size cell number x ensemble number, we determine the enlistment of neurons into individual ensembles by only including neurons whose entry is over 1 standard deviation from the mean for their ensemble. Only detected ensembles that were comprised of at least four neurons were included in further analysis. When analyzing spontaneous recordings, ensembles were defined from the raster of the entire recording. In experiments where visual stimuli were presented to animals, ensembles were defined only from the raster during the stimulus period.

### Quantifying ensemble expression and a threshold for ensemble detection

Here we define ensemble expression as the percent of an ensemble’s neurons synchronously active at every frame. For each ensemble, the overall raster of neuronal activity was constrained to only the neurons comprising that ensemble, and then ensemble expression was quantified as the percent of cellular activation across these neurons at each frame. Combining the activity from all of our recordings, we observed a long tail on the distribution of ensemble expression beginning 3 standard deviations from the mean (35%; Fig. S1A-B), and we defined this value as the threshold for ensemble activation. From this we labeled an ensemble as active at a given frame if the ensemble expression was greater than 35%. The analysis of ensemble expression only included values over threshold.

### Ensemble specificity

Ensembles were separated using Pearson’s linear correlation (α = 0.01) into three major categories: stimulus-specific, locomotion-specific, and non-sensorimotor. An ensemble was defined as stimulus-specific if the ensemble expression had a significant positive correlation with a vector denoting when stimuli presentations occurred. An ensemble was defined as locomotion-specific if the ensemble expression was positively associated with a vector labeling locomotion. Non-sensorimotor ensembles were all other ensembles not labeled as related to stimulus or locomotion. Further segregation of non-sensorimotor ensembles was performed into three categories: anticorrelated to stimulus, sensory trending, and correlated to delta activity. Ensembles anticorrelated were ensembles that were significantly negatively correlated to stimulus presentation. Sensory trending ensembles were not correlated to overall stimulus presentation and correlated to the presentation of only one orientation of stimulus. Non-sensorimotor delta related ensembles were positively correlated to delta.

## Statistical Analysis

All data points represent the average of that specific variable over an entire individual imaging session. Statistical testing on the same variable in time was performed using nonparametric statistical testing with an alpha of 0.05. On the comparison of variables across brain state, if a significant effect from the Kruskal-Wallis test was found using an alpha of 0.05, then further testing was performed across individual brain states using Dunn’s multiple comparison tests. On the comparison of variables across brain state before and after visual stimulus presentation, a Friedman test was performed, and a post hoc Wilcoxon test was performed on each brain state before vs. after visual stimulus presentation with a Bonferroni correction against an alpha of 0.05.

## Acknowledgements

We would like to thank Lux Steinberg for technical support and help with writing and members of the Yuste lab; for helpful discussions; This work was supported by NIH-NIA (F32AG354069) and (K99AG084853) to J.L.; and NEI (R01EY035248) to R.Y.

## Author Contributions

J.L. and R.Y. contributed to project conception, project design, and manuscript writing. J.L. performed the experiments and analyzed the results. J.L. and R.Y. directed the project and secured resources and funding.

## Competing Interests Statement

The authors declare no competing interests.

## Supporting Information

**Fig. S1.**
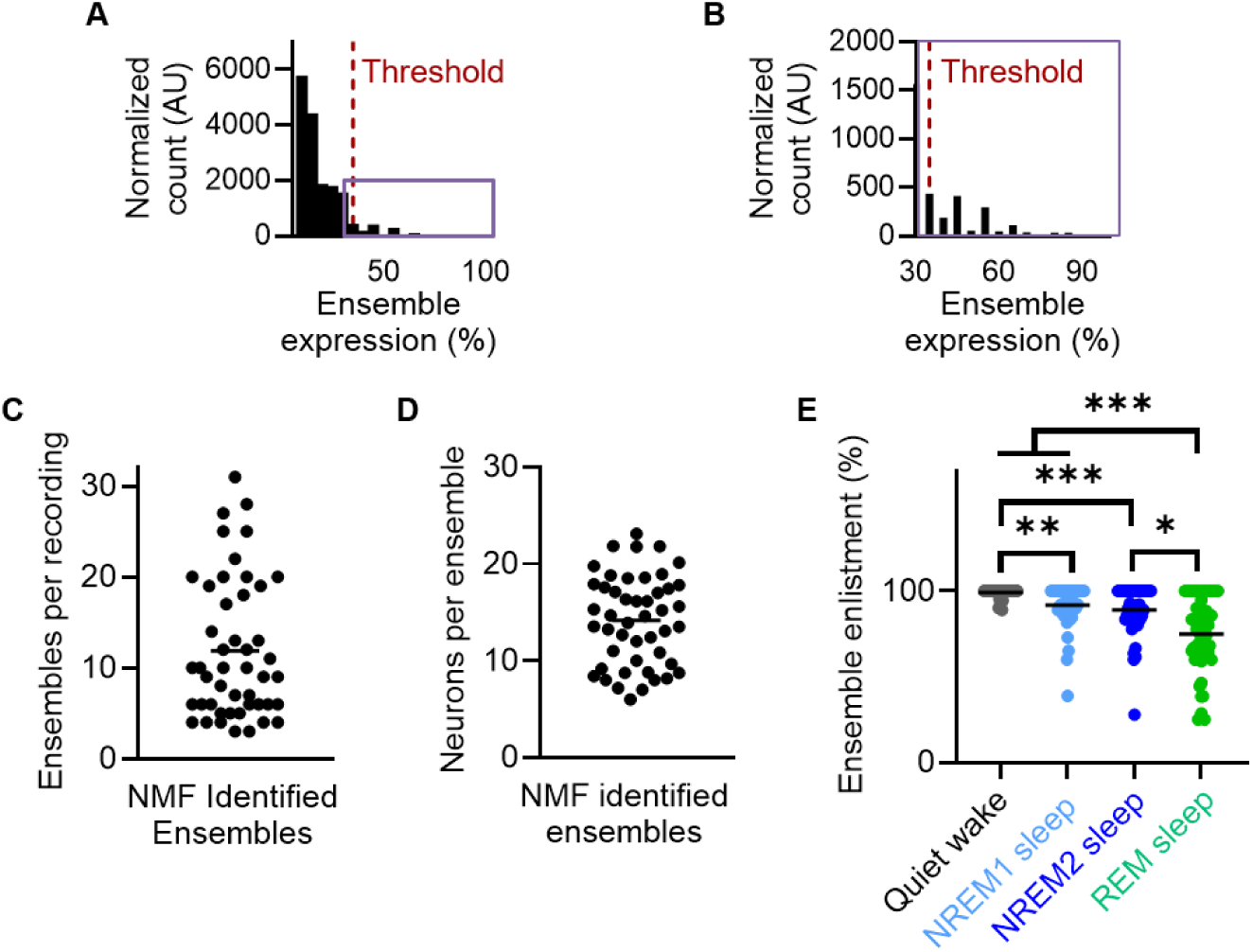
Characteristics of neuronal ensembles in visual cortex across brain states. (**A**) Histogram of counts of ensemble expression across all frames, across all recordings. Note, the red line depicting three standard deviations from the mean (35.0%) ensemble threshold at the start of the long tail in the distribution. Lavender box denotes range of B. (**B**) Same as A, but for suprathreshold values for ensemble expression. (**C**) Number of ensembles detected using NMF across spontaneous recordings. (**D**) Number of neurons per ensembles across spontaneous recordings. (**E**) The percent of ensembles present across different brain states for each recording. Dunn’s test after significant Kruskal-Wallis test. * ≡ p<0.05, *** ≡ p<0.01, *** ≡ p<0.001

**Fig. S2.**
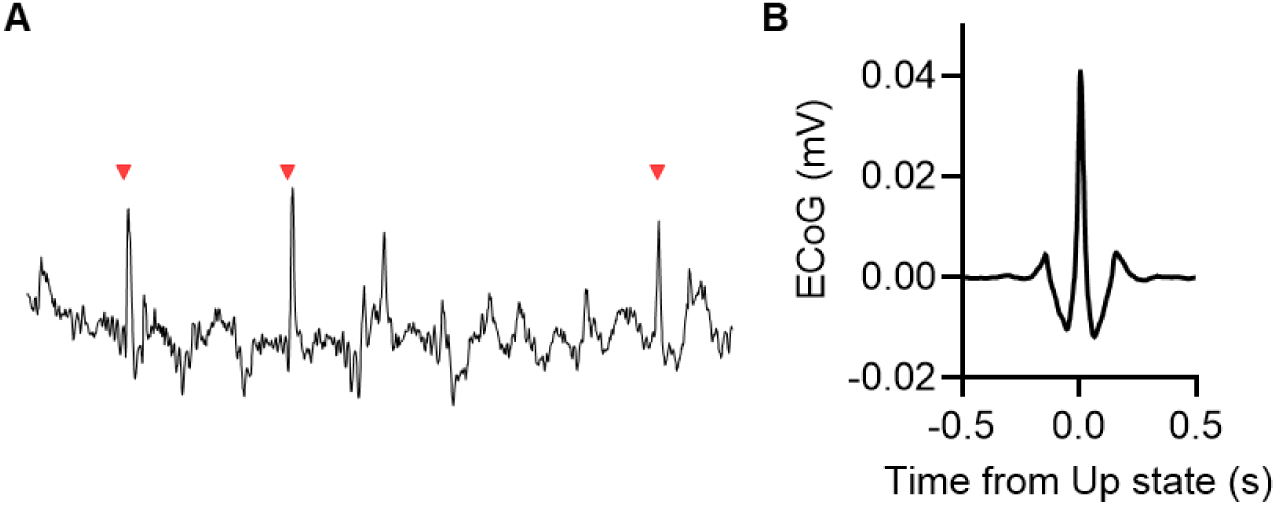
Detecting cortical UP states during NREM sleep. (**A**) Example detected UP states (red arrows) during NREM sleep. (**B**) Average Up state across recordings when animals were in NREM sleep.

**Fig. S3.**
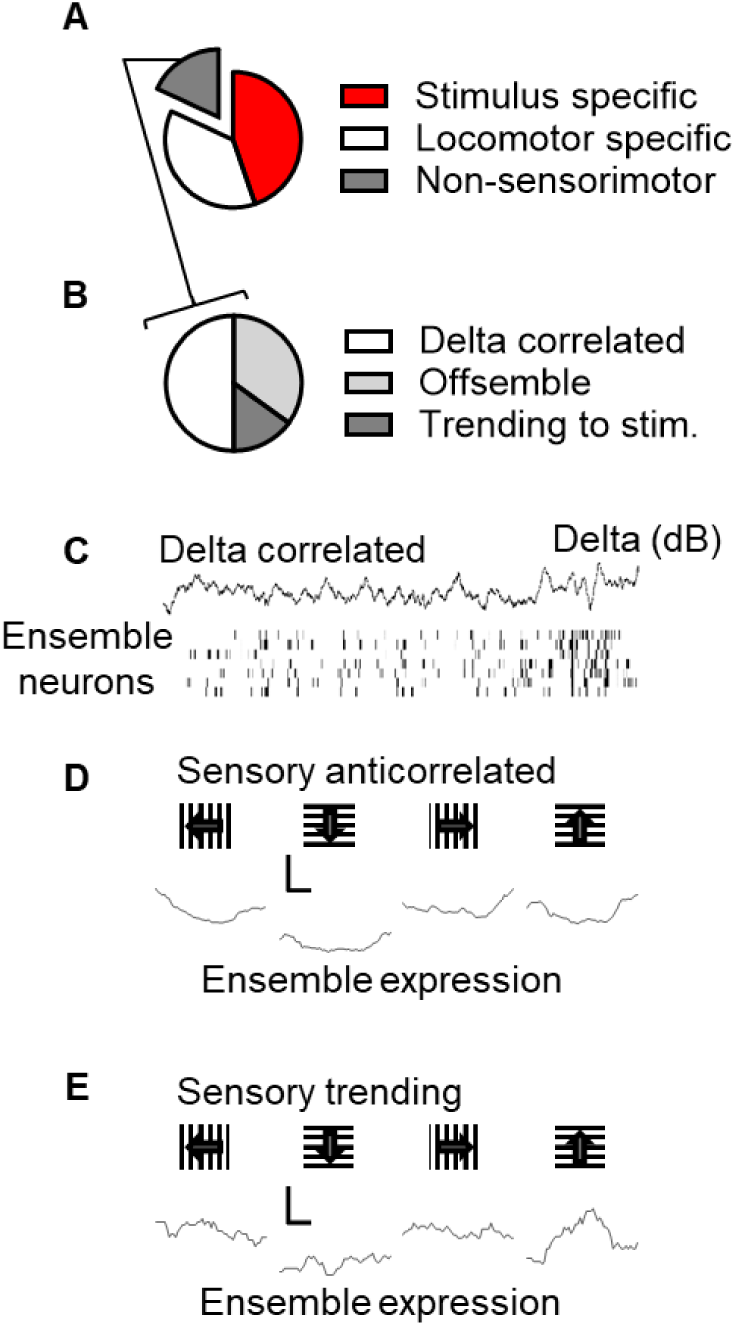
Ensemble categorization. (**A**) Proportion of ensemble classes found across all ensembles found (n=286). (**B**) Makeup of non-sensorimotor ensemble classes found (n=52). (**C**) Example of an ensemble correlated to delta power during the stimulus period. (**D**) Example of an ensemble that is anticorrelated to sensory stimulation. (**E**) Example of an ensemble trending to stimulus response.

## Notes

### Competing Interest Statement

The authors have declared no competing interest.

### Summary of Updates

Sample sizes have been increased and additional analysis provided to uncover the cellular activity during hypnagogic sleep states as well as data suggesting cortical neuronal ensembles are formed from a pre-allocated lexicon. The addition of Figures S2-4 also add additional information related to ensemble labeling, ECoG UP-state analysis, and ensemble-specific encoding.

